# ATP1A3 dysfunction alters excitability and developmental trajectory of human neurons in alternating hemiplegia of childhood

**DOI:** 10.64898/2026.10.09.757992

**Authors:** Christopher H. Thompson, Christine Q. Simmons, Syed M.A. Wafa, Alfred L. George

**Author notes:** contributed equally (Christopher H. Thompson, Christine Q. Simmons).

## Abstract

Pathogenic variants in *ATP1A3*, encoding the neuronal Na/K-ATPase, are associated with a spectrum of neurodevelopmental disorders including alternating hemiplegia of childhood (AHC). *ATP1A3* is expressed throughout the cortex during early development, and later becomes enriched in GABAergic neurons. Here, we interrogated the underlying mechanisms and neurophysiological consequences of a recurrent AHC-associated variant (*ATP1A3*-G947R) in human induced pluripotent stem cell (hiPSC) derived glutamatergic and GABAergic neurons. We found that glutamatergic and GABAergic AHC neurons exhibited smaller ouabain-sensitive pump currents compared to isogenic mutation-corrected control neurons. This impairment of pump function correlated with a depolarized potassium reversal potential, lower intracellular potassium concentration, and depolarized resting membrane potential for both cell types compared to isogenic control neurons. Additionally, using a Na^+^ sensitive dye, we observed that AHC neurons had a higher basal intracellular Na^+^ concentration that recovered more slowly following Na^+^ loading compared to isogenic control neurons. As an additional consequence of impaired pump function and abnormal Na^+^ dynamics, glutamatergic and GABAergic AHC neurons were more prone to depolarization block. Finally, the time course by which GABA neurotransmission switches from excitatory to inhibitory was delayed in glutamatergic AHC neurons compared to isogenic controls. Our findings provide evidence that impairment of ATP1A3 function has direct effects on ionic homeostasis, leading to cascading effects on intrinsic neuronal firing and maturation of GABAergic neurotransmission that advance understanding of the pathophysiology of AHC.

## Introduction

Heterozygous pathogenic variants in *ATP1A3*, which encodes the catalytic (α3) subunit of the neuronal Na/K-ATPase, are associated with a spectrum of neurodevelopmental disorders including alternating hemiplegia of childhood (AHC) (1–7). Individuals affected by AHC typically experience transient attacks of one-sided or bilateral weakness and/or dystonia with varying duration, abnormal eye movements, seizures, hypotonia, disordered sleep, and delayed neurodevelopment (8–11), and generally poor long-term neurological outcomes often with persistent motor dysfunction, impaired communication, and cognitive disabilities (12). Life-threatening events including apnea and sudden unexpected death may occur in AHC (13–15). There are no disease-modifying treatments available and symptom management is standard of care (16). While much has been learned about the heterogeneous clinical features, natural history and genetic basis of AHC since 2012 (17), the cellular consequences of *ATP1A3* dysfunction in human neurons and fundamental pathophysiological mechanisms that underlie this disorder remain poorly understood.

The Na/K-ATPase (NKA) is a catalytic ion pump essential for establishing ion gradients and maintaining electrical stability in cells. By coupling the hydrolysis of ATP with the export of three Na^+^ ions and import of two K^+^ ions, the NKA generates asymmetric transmembrane ion concentration gradients that determine the resting membrane potential and enable the generation of action potentials in electrically excitable cells. In neurons, the *ATP1A3*-encoded NKA is particularly important for rapidly extruding Na^+^ and restoring intracellular Na^+^ levels after intense neuronal activity, when local intracellular Na^+^ concentrations can increase transiently by 30-100 mM (18, 19). *ATP1A3* is expressed beginning in early human brain development and persists throughout postnatal life (20), which helps explain the severity of neurodevelopmental disorders associated with pathogenic variants in this gene.

Three recurrent and often *de novo ATP1A3* variants (D801N, E815K, and G947R) account for approximately 60% of AHC cases. While D801N and E815K are the most prevalent variants in large AHC cohorts including participants mostly from Europe and the United States (21, 22), G947R was recently recognized as the most frequent *ATP1A3* variant in a large case series from Brazil (23). Two recurrent variants (D801N, G947R) affect residues within or adjacent to known cation binding sites (24). We previously reported preliminary evidence that human glutamatergic neurons heterozygous for the AHC-associated recurrent variant *ATP1A3*-G947R exhibited smaller ouabain-sensitive pump current, which is a direct measure of ion transport, and a collapsed K^+^ gradient that correlated with a depolarized resting membrane potential (25). An important limitation of this work was that *ATP1A3*-G947R neurons were compared to healthy control neurons, which does not control for the influence of genomic background. In addition, the impact of ATP1A3 dysfunction on other neuron subtypes and the impact of pump dysfunction on intracellular Na^+^ dynamics were not investigated.

Here we present comparisons of *ATP1A3*-G947R neurons with isogenic mutation-corrected neurons, which better assess the pathogenic effects of this variant without potential confounding factors contributed by different genomes. We studied both glutamatergic and GABAergic neurons to assess the impact of this mutation on properties ranging from ionic gradients, intracellular Na^+^ dynamics, neuronal physiology, and neurodevelopmental trajectory. Our findings establish important features of the cellular pathophysiology of AHC helping to advance our understanding of the fundamental mechanisms responsible for paroxysmal neurological features and impaired neurodevelopment in this disorder.

## Results

### Generation of isogenic mutation-corrected iPSC line

We used genome editing based on clustered regularly interspaced short palindromic repeats (CRISPR) to correct the *ATP1A3*-G947R variant in a patient-derived iPSC line (25). Additional synonymous single nucleotide variants (SNV) were intentionally generated in the protospacer adjacent motif (PAM; c.2829C>T) and the third base position of the corrected glycine codon (c.2841G>A) to limited DNA re-cleavage and to help exclude hemizygosity, respectively (**Fig. 1A**). Several iPSC clones were sequenced to identify lines with the intended edits, to exclude on-target editing errors such as insertions or deletions, and to exclude predicted off-target edits elsewhere in the genome. Then sequence-correct clones were karyotyped to exclude large chromosomal alterations (**Supplemental Figure S1**). One clonal line that satisfied all of these quality control tests (**Fig. 1B**) was selected for use as an isogenic mutation-corrected control.

**Figure 1.**
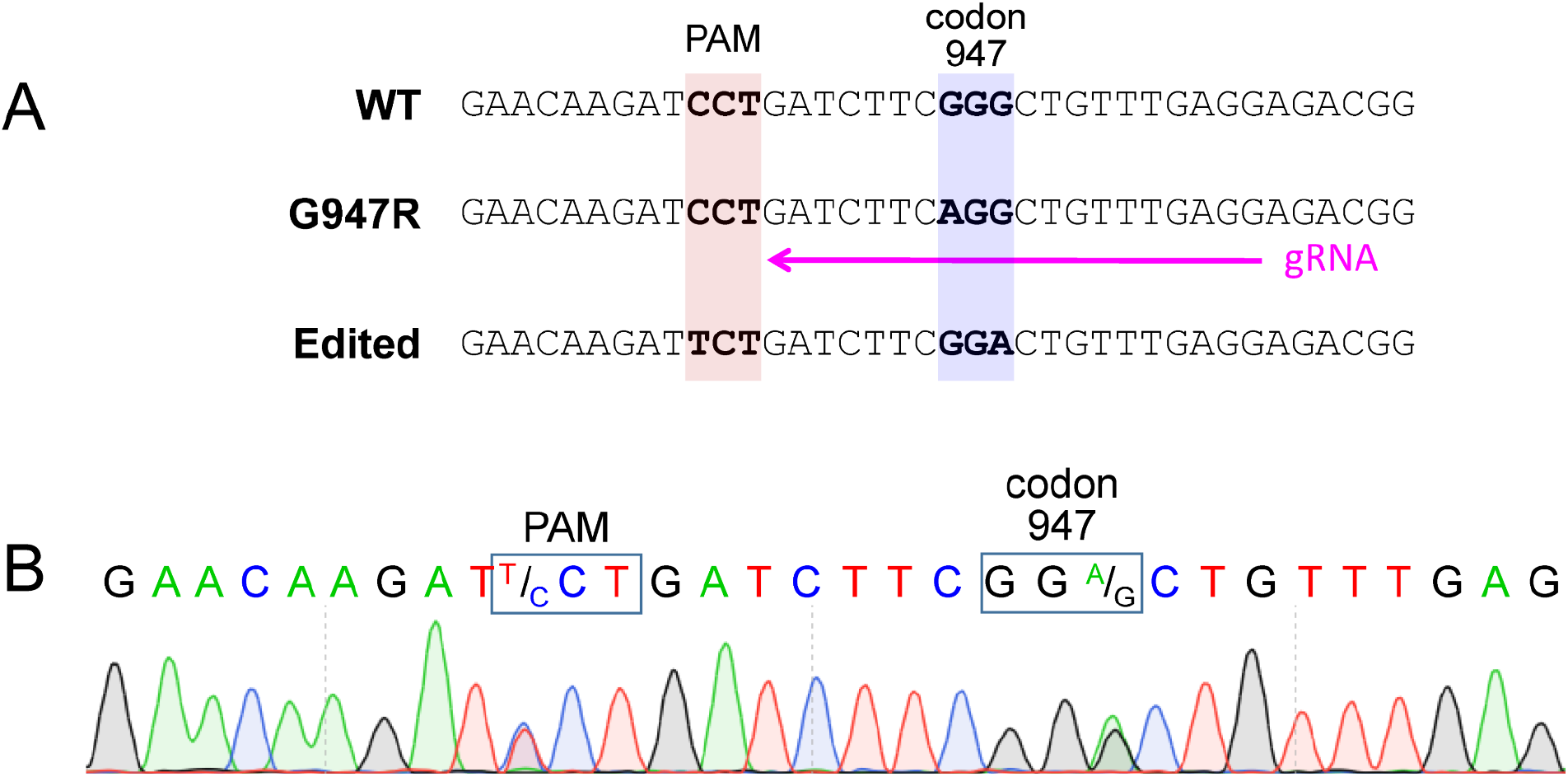
Design and validation of mutation-corrected isogenic iPSC line. (**A**) Design of CRISPR genome editing of *ATP1A3*-G947R. Positions of the PAM site and guide RNA (gRNA) refer to the antisense strand. (**B**) Sanger sequence trace illustrating genome edited *ATP1A3* sequence (nucleotide 2826-2852 based on NCBI RefSeq NM_152296). Boxed nucleotides represent edited PAM site with heterozygous synonymous SNV (C>T), and edited codon-947 with heterozygous synonymous SNV (G>A).

Glutamatergic and GABAergic neurons were differentiated in parallel from the heterozygous A*TP1A3*-G947R and isogenic control lines then co-cultured with primary mouse cortical glial cells. GABAergic neurons were plated with glutamatergic neurons in a 30:70 ratio to enhance long-term viability and promote maturation. The different neuron subtypes in co-cultures were distinguished by lentivirus-encoded fluorescent proteins introduced at the time of differentiation. Data from 834 cells were included in this study. All measurements and statistical analyses were conducted blind to genotype.

### AHC neurons exhibit smaller pump currents

To directly assess ion transport by Na/K ATPase in human neurons, we measured total and ouabain-sensitive pump current, which is generated by the net outward movement of positive charge coincident with Na^+^ efflux. Upon exposure to 20 mM extracellular K^+^, heterozygous *ATP1A3*-G947R glutamatergic and GABAergic neurons exhibited smaller total and ouabain-sensitive pump current than isogenic control neurons **(Fig. 2; Table S1)**. There were no differences in baseline current recorded in the absence of extracellular K^+^ either in the presence or absence of ouabain (**Supplemental Fig. S2**; **Table S1**). These data are consistent with impaired ion transport caused by this AHC-associated pathogenic variant.

**Figure 2.**
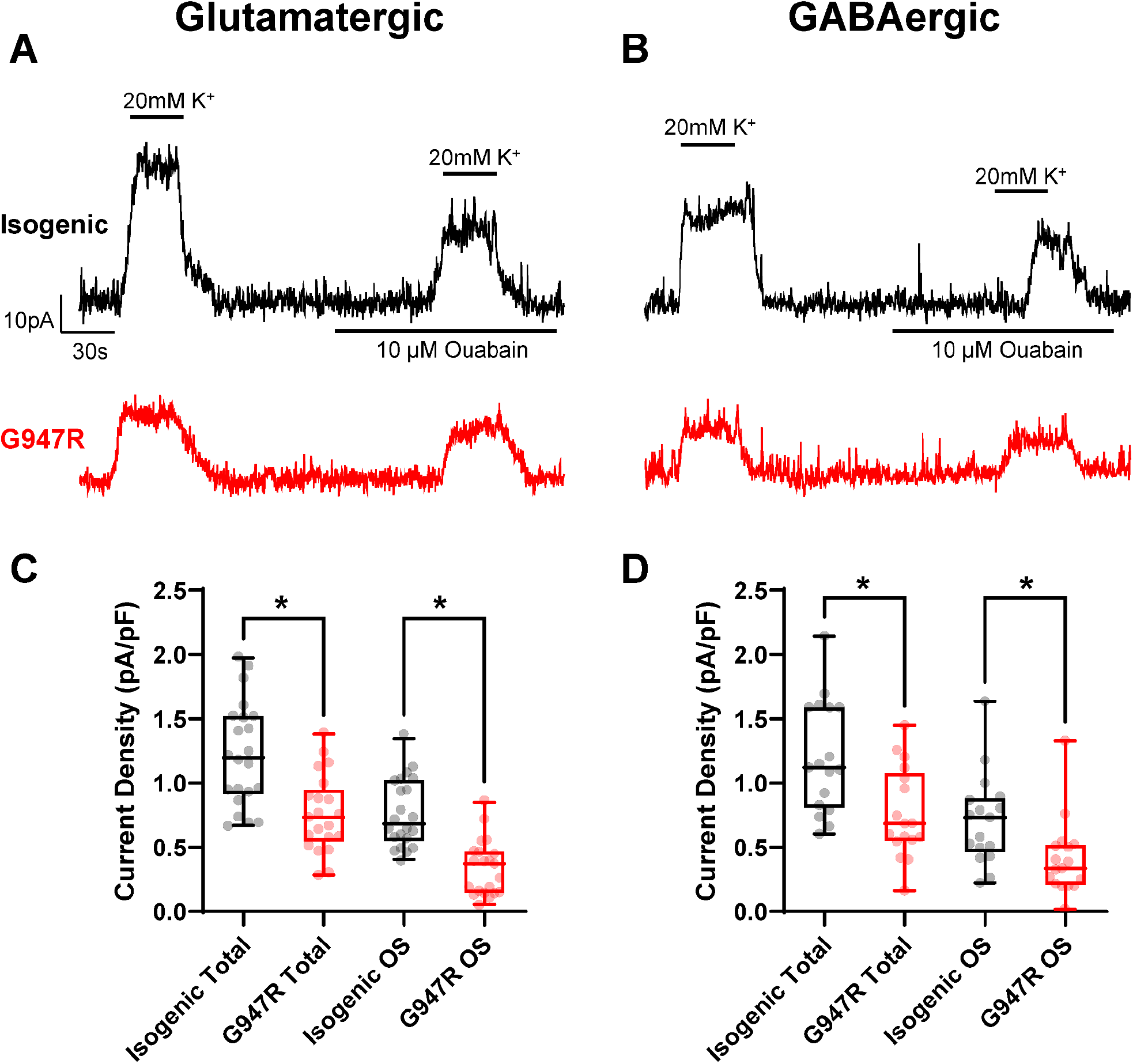
ATP1A3-G947R exhibit smaller pump currents. Representative current traces of outward pump current activated by 20 mM K^+^ in the absence and presence of 10 µM ouabain in **(A)** glutamatergic and **(B)** GABAergic iPSC-derived neurons for isogenic control (black) and G947R (red) neurons. Summary data for total and ouabain sensitive (OS) pump current for **(C)** glutamatergic and **(D)** GABAergic neurons. Box plots represent median ± 95% CI for 17 – 22 cells. Asterisks denote p<0.05.

### AHC neurons exhibit a collapsed K^+^ gradient and depolarized resting potential

Impaired Na/K-ATPase ion transport may affect steady-state ion concentration gradients. To investigate the transmembrane K^+^ gradient in *ATP1A3*-G947R neurons we measured the potassium equilibrium potential (*E*_K_) and then used these values to infer the intracellular K^+^ concentration. We conducted experiments with isogenic pairs of heterozygous G947R and isogenic control glutamatergic or GABAergic neurons. We found that *E*_K_ was significantly depolarized in both types of heterozygous mutant neurons compared to isogenic control cells **(Fig. 3A, B; Table S2)**. Using the measured values of *E*_K_, we calculated the intracellular K^+^ concentration with the Nernst equation. This calculation predicted that heterozygous mutant glutamatergic and GABAergic neurons have significantly lower intracellular K^+^ concentrations than isogenic control cells **(Fig. 3C, D; Table S2)**.

**Figure 3.**
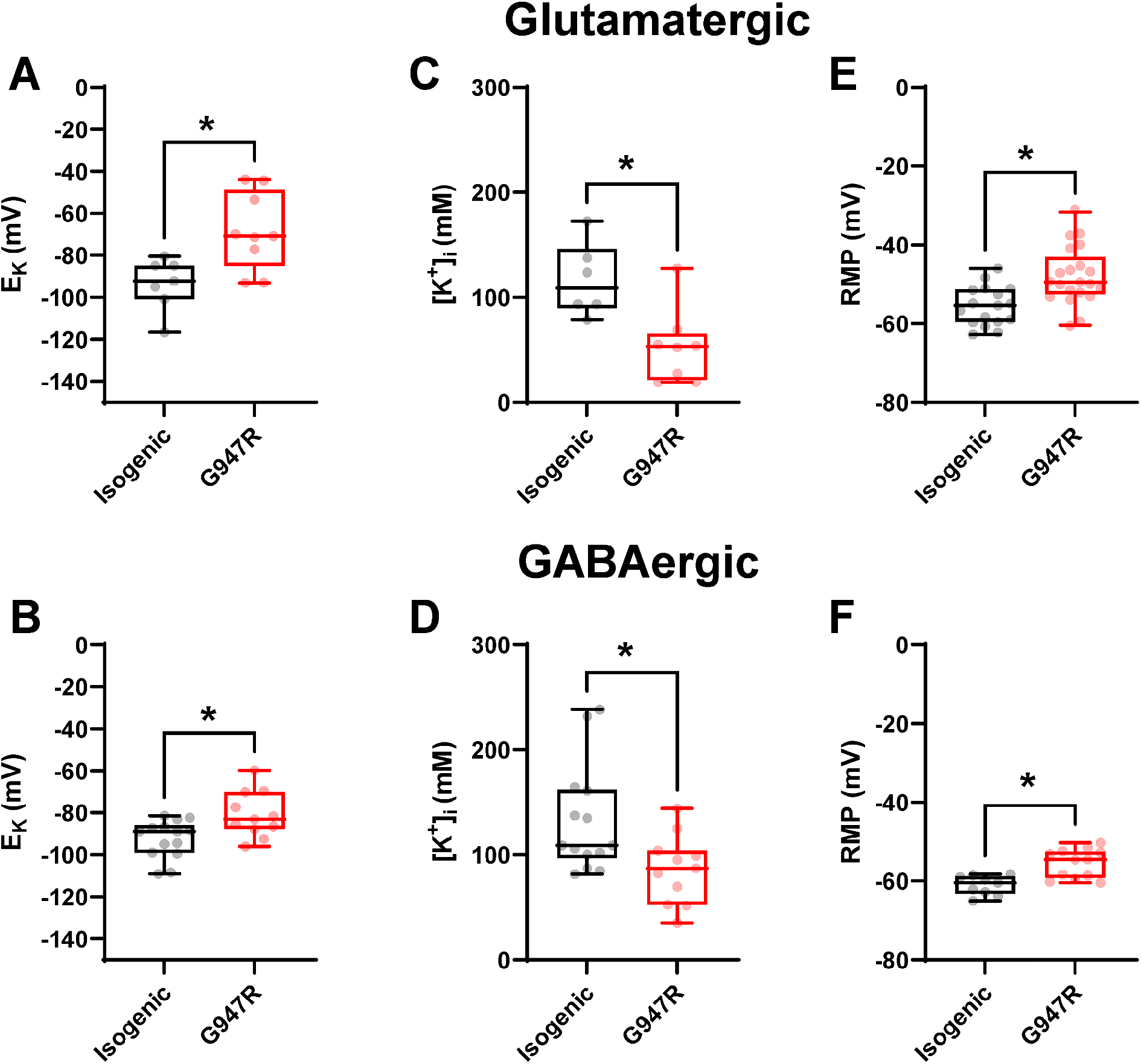
ATP1A3-G947R have altered resting membrane potential and K^+^ equilibrium potential. Summary data for K^+^ equilibrium potential (E_K_) in (**A**) glutamatergic neurons, and (**B**) GABAergic neurons, calculated intracellular K^+^ concentration in (**C**) glutamatergic neurons and (**D**) GABAergic neurons, and resting membrane potential (RMP) in (**E**) glutamatergic neurons and (**F**) GABAergic neurons. (Box plots represent median ± 95% CI for 6 - 21 cells. Asterisks denote p<0.05.

From these observations, we predicted that an attenuated transmembrane K^+^ gradient will blunt the chemical driving force on K^+^ leak channels that contribute to setting the resting membrane potential (RMP). Using the intracellular K^+^ concentrations inferred from the measured *E*_K_ values, we used the Goldman-Hodgkin-Katz equation to predict the theoretical RMP in the neurons. The predicted RMP for isogenic control glutamatergic neurons was −58.7 ± 3.7 mV compared to −45.9 ± 4.6 mV for heterozygous G947R neurons. Likewise, predicted RMP for isogenic control GABAergic neurons was −60.4 ± 3.1 mV compared to −54.0 ± 3.2 mV for G947R neurons. To test these predictions, we measured RMP directly using perforated patch current clamp recording. The measured RMP was significantly depolarized for both glutamatergic and GABAergic G947R neurons compared to isogenic control cells **(Fig. 3E, F; Table S2)**. The close alignment of measured RMP with that predicted based on calculated intracellular K^+^ concentrations supports our contention that disruption of the K^+^ concentration gradient is a major driver of the depolarized RMP.

### AHC neurons exhibit altered intracellular Na^+^ dynamics

We reasoned that impaired pump function will also impact intracellular Na^+^ concentration. To test this, we used a Na^+^ sensitive dye (ING-2) to measure basal intracellular Na^+^ concentration in isogenic control and heterozygous G947R glutamatergic neurons. Baseline fluorescence intensity for G947R neurons was higher compared to isogenic control neurons **(Fig. 4A; Table S2)**. Because ING-2 intensity directly correlates with Na^+^ concentration, this indicates that G947R neurons have a higher basal intracellular Na^+^ concentration than isogenic control neurons.

**Figure 4.**
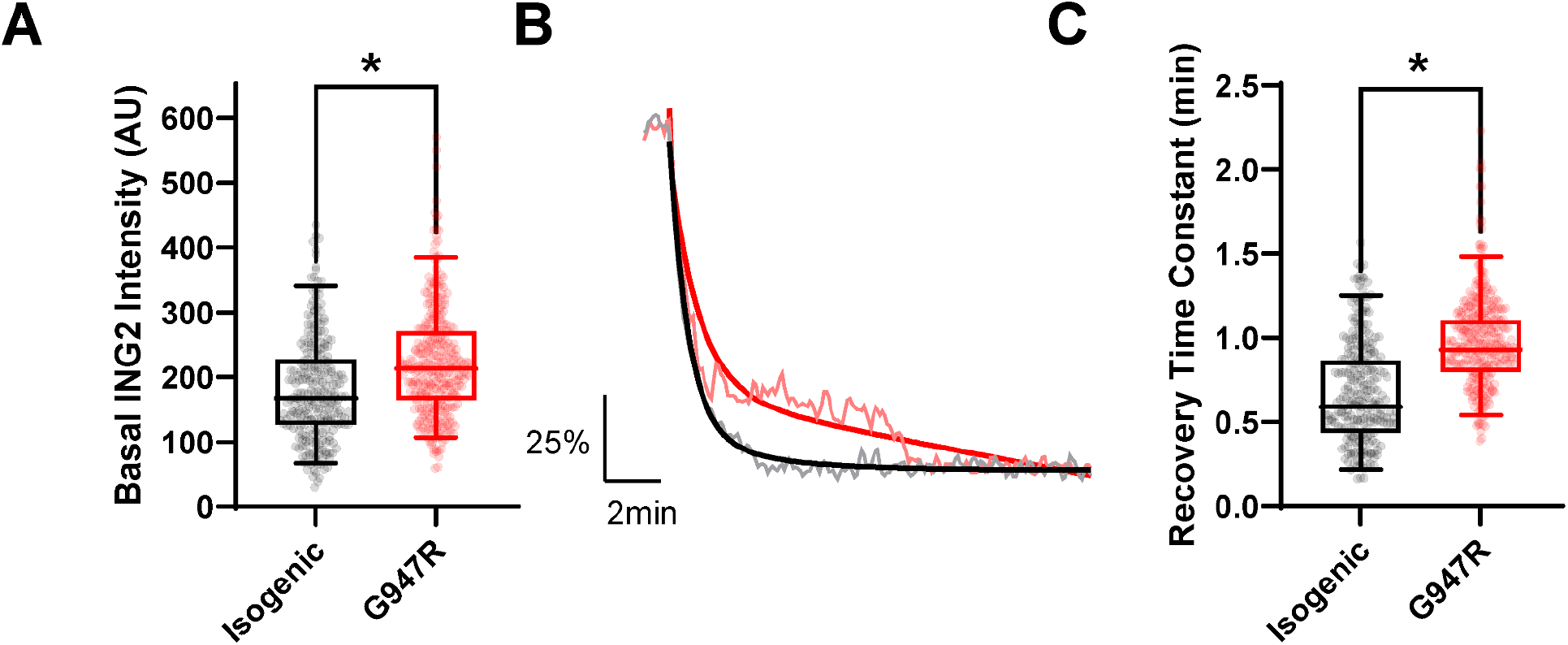
ATP1A3-G947R neurons exhibit altered intracellular Na^+^ dynamics. **(A)** Summary data showing basal ING-2 fluorescence intensity (A.U.) in isogenic control (black) and ATP1A3-G947R (red) NGN2 glutamatergic neurons. **(B)** Average normalized fluorescence intensity traces for isogenic control (black) and ATP1A3-G947R (red) glutamatergic neurons highlighting recovery of basal sodium concentration following Na^+^ loading. Solid smooth lines represent the average exponential fit of the decay phase. **(C)**. Summary data for the time constant for recovery following Na^+^ loading for isogenic control (black) and ATP1A3-G947R (red) glutamatergic neurons. Box plots represent median ± 95% Ci for 250 – 321 cells. Asterisks denote p<0.05.

Another important function of ATP1A3 is to clear accumulated intracellular Na^+^ following high frequency firing. To investigate if this function was affected by the G947R mutation, we loaded neurons with high Na^+^ by continuous exposure of cells to a high Na^+^ extracellular solution after dye loading. We then measured the recovery time constant following application of an extracellular solution containing low (10 mM) Na^+^ concentration. The time constant for recovery from Na^+^ loading was significantly larger for G947R glutamatergic neurons compared to isogenic control neurons indicating a slower recovery time course **(Fig. 4B,C; Table S2)**. The sparse distribution of GABAergic neurons in co-culture with glutamatergic neurons precluded us from measuring sodium dynamics with the desired throughput. These results suggest that AHC neurons maintain a blunted transmembrane Na^+^ concentration gradient and have impaired capacity for Na^+^ clearance.

### AHC neurons are more prone to depolarization block

We investigated the impact that the blunted transmembrane Na^+^ gradient and impaired Na^+^ clearance may have on neuronal excitability of AHC neurons. Using perforated patch current clamp to preserve transmembrane ionic gradients, we measured action potential generation during 2-second graded current stimulations. Glutamatergic G947R neurons exhibited substantially lower firing frequency at strong stimuli compared to isogenic control neurons and had a greater tendency for depolarization block **(Fig. 5A)**. We quantified the time to depolarization block when repetitive firing ceased. Mutant glutamatergic neurons entered depolarization block more quickly than isogenic control neurons **(Fig. 5B,C; Table S3)**. We also examined neuronal excitability and time to depolarization block in GABAergic neurons and found that while overall neuronal excitability was similar between isogenic control and G947R neurons, mutant GABAergic neurons exhibited depolarization block more quickly **(Fig. 5C,D,E; Table S3)**.

**Figure 5.**
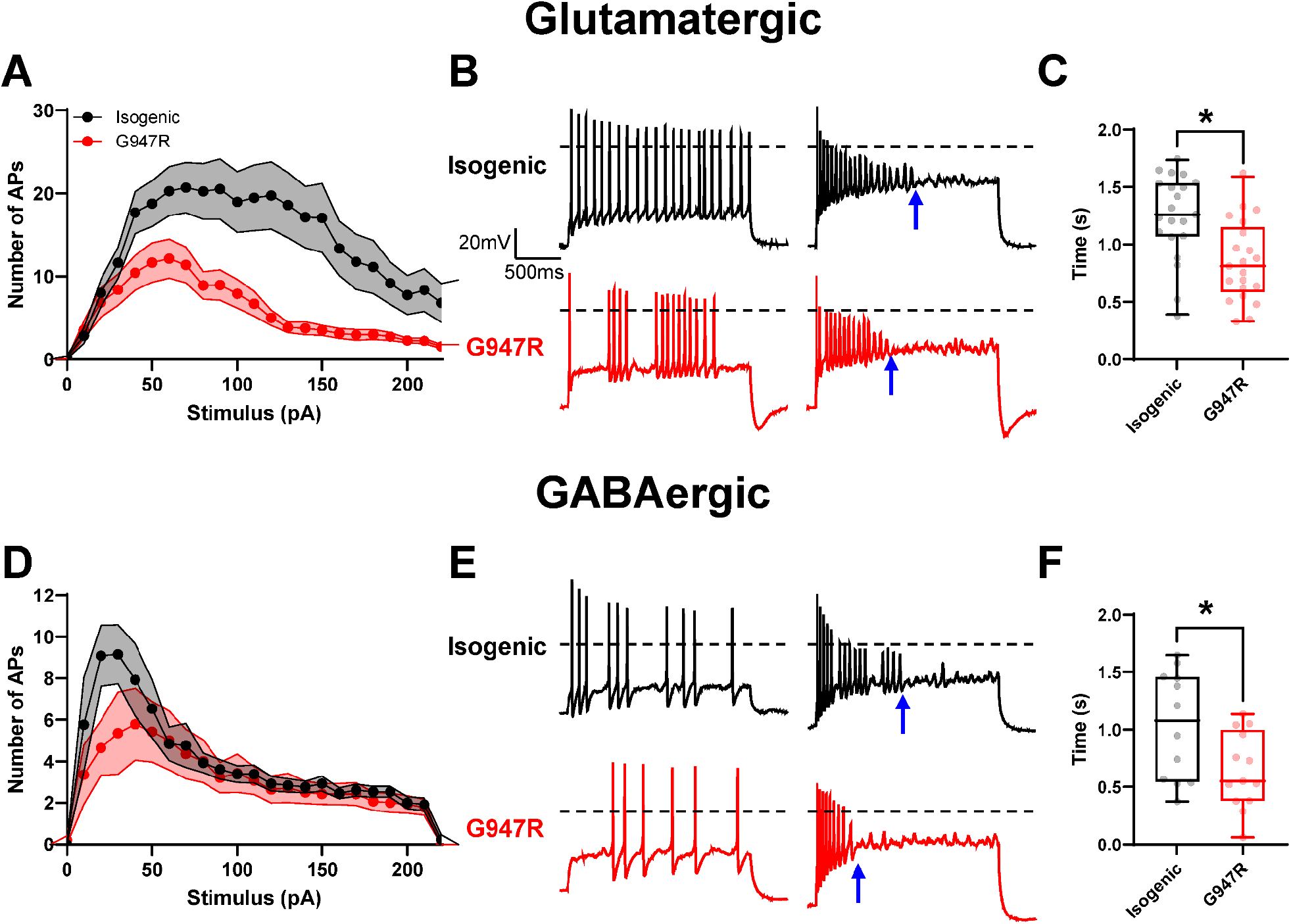
ATP1A3-G947R neurons are more prone to depolarization block elicited by step currents. **(A)** Summary input-output curves for isogenic control (black) and G947R (red) glutamatergic neurons. **(B)** Representative traces showing repetitive neuronal firing and onset of depolarization block (blue arrow) recorded from isogenic control (black) and G947R (red) glutamatergic neurons. **(C)** Summary data showing time required to induce depolarization block in glutamatergic neurons. **(D)** Summary input-output curves for isogenic control (black) and G947R (red) GABAergic neurons. **(E)** Representative traces showing repetitive neuronal firing and onset of depolarization block (blue arrow) recorded from isogenic control (black) and G947R (red) GABAergic neurons. **(F)** Summary data showing time required to induce depolarization block in GABAergic neurons. Input-output curves are plotted as mean ± SEM and box plots represent median ± 95% CI from 12 - 21 cells. Asterisks denote p<0.05.

To further quantify how susceptible neurons are to depolarization block, we used a 3-second ramp protocol to quantify the absolute number of action potentials fired before onset of depolarization block, and the membrane potential at which depolarization block occurred. We found that isogenic control glutamatergic neurons fired nearly twice as many action potentials before exhibiting depolarization block compared to G947R neurons **(Fig. 6A,B; Table S3)**. Additionally, isogenic control neurons were able to reach more depolarized potentials before the onset of depolarization block **(Fig. 6A,C; Table S3)**. Finally, G947R neurons exhibited depolarization block with a weaker stimulus, compared to isogenic control neurons (**Supplemental Fig. S3**; **Table S3**). We observed similar trends for GABAergic neurons, where isogenic control neurons were able to fire more action potentials **(Fig. 6D,E; Table S3)** and reach a more depolarized membrane potential **(Fig. 6D,F; Table S3)**. These results suggest that both mutant glutamatergic and GABAergic neurons are more prone to depolarization block, which we contend is related to impaired Na^+^ clearance.

**Figure 6.**
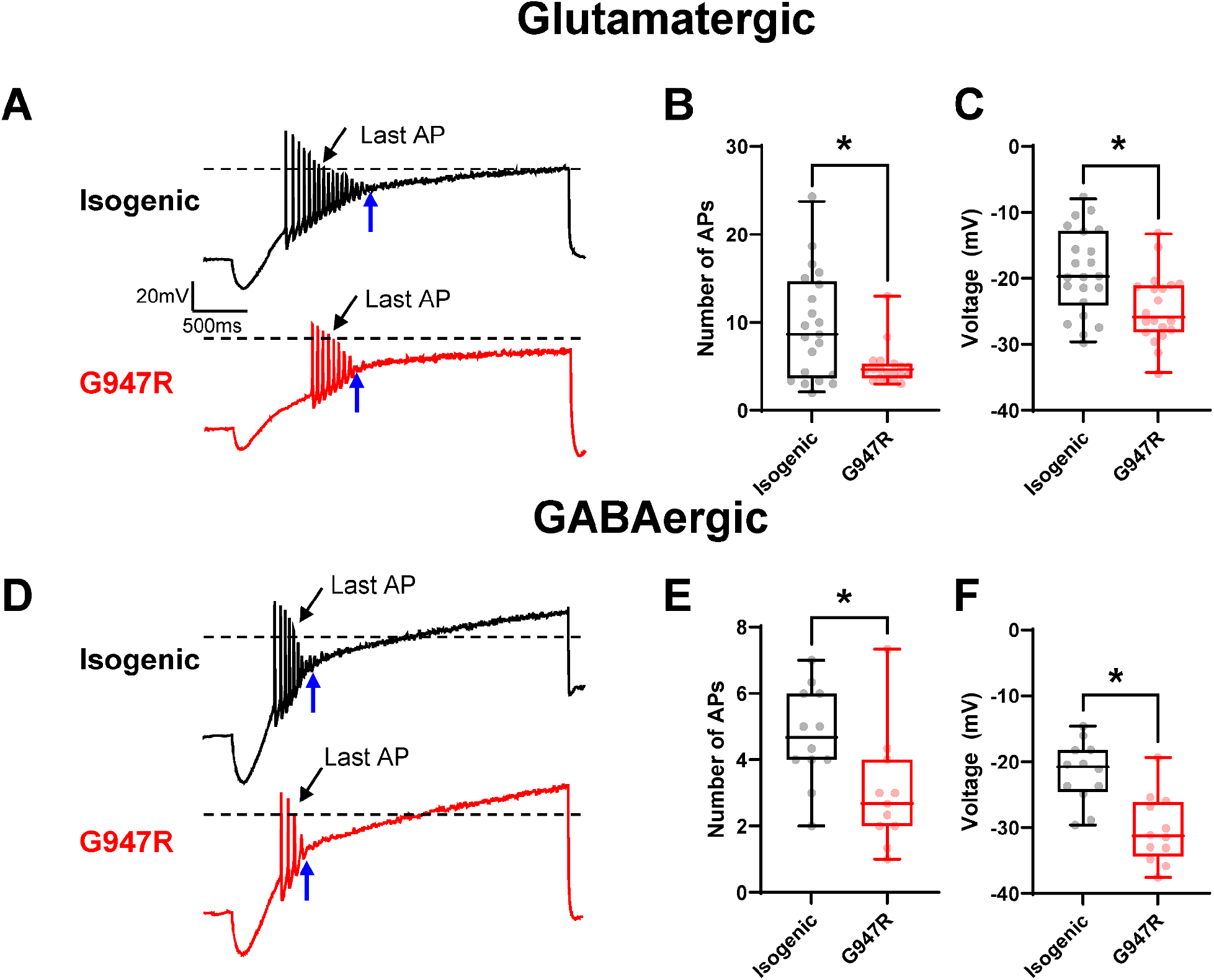
ATP1A3-G947R neurons are more prone to depolarization block elicited by ramp currents. **(A)** Representative traces of action potentials evoked by ramp currents recorded from isogenic control (black) and G947R (red) glutamatergic neurons. Onset of depolarization block occurs at the blue arrows. Summary data showing **(B)** the number of action potentials fired before induction of depolarization block, and **(C)** the membrane potential at the onset of depolarization block in glutamatergic neurons. **(D)** Representative traces of action potentials evoked by ramp currents recorded from isogenic control (black) and G947R (red) GABAergic neurons. Onset of depolarization block occurs at the blue arrows. Summary data showing **(E)** the number of action potentials fired before induction of depolarization block, and **(F)** the membrane potential at the onset of depolarization block in GABAergic neurons. Box plots represent median ± 95% CI for 11 – 22 cells. Asterisks denote p<0.05.

### AHC neurons exhibit differences in action potential morphology

Action potential morphology is shaped by underlying ionic conductances as well as how neurons manage the ionic driving forces during a train of action potentials. We examined action potential morphology of isogenic pairs of glutamatergic and GABAergic neurons to determine how morphological differences that emerge during the course of successive action potentials contribute to the onset of depolarization block. For glutamatergic neurons, we found that the first action potential elicited in a train of action potentials was largely similar between mutant and isogenic control neurons. Only action potential amplitude) and the fast component of the afterhyperpolarization (fAHP) were different between genotypes **(Fig. 7; Table S4)**. However, when comparing the last action potential in a train, several differences were apparent. Compared to isogenic control neurons, the last action potential elicited for G947R neurons had a more depolarized threshold, smaller amplitude, wider half-width, which were associated with slower upstroke and downstroke velocities, and less hyperpolarized fAHP **(Fig. 7D-G; Supplemental Fig. S4; Table S4)**.

**Figure 7.**
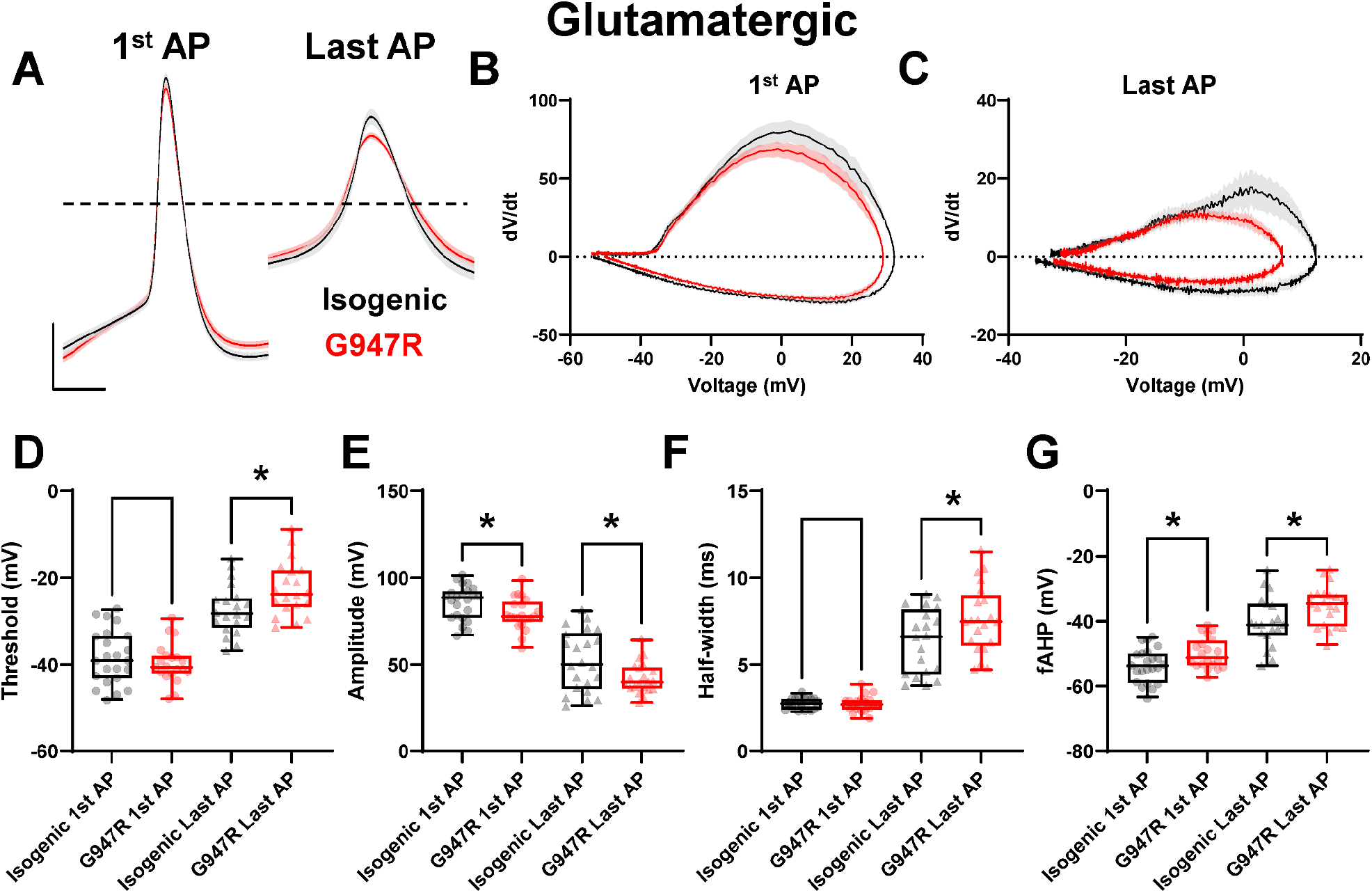
Action potential morphology of glutamatergic neurons. **(A)** Averaged first and last action potentials evoked by 100 pA current inputs for isogenic control (black) and G947R (red) glutamatergic neurons. Averaged phase plane plots for the first **(B)** and last **(C)** action potentials induced by 100 pA recorded from isogenic control (black) and G947R (red) glutamatergic neurons. Summary data for **(D)** action potential threshold, **(E)** amplitude, **(F)** half-width, and **(G)** fAHP for the first (circles) and last (triangles)action potentials evoked by 100 pA current inputs. Average action potentials and phase plane plots are plotted as mean ± SEM and box plots represent median ± 95% CI for 19 – 23 cells. Asterisks denote p<0.05.

Action potential morphology was similar between G947R and isogenic control GABAergic neurons, regardless of whether the action potential was the first or last action potential in a train **(Fig. 8; Supplemental Fig. S4; Table S5)**. However, upstroke and downstroke velocity **(Table S5)** were faster for the last action potentials in a train for G947R neurons compared to isogenic control neurons **(Fig. 8B,C; Table S5)**. Additionally, the last action potential in the train for GABAergic G947R neurons had larger amplitude compared to isogenic mutation-corrected neurons **(Fig. 8E; Table S5)**. Input resistance was not different between isogenic and G947R neurons for either cell type (**Supplemental Fig. S5; Tables S4 and S5**) Thus, for glutamatergic neurons, differences in action potential morphology are more pronounced in G947R neurons, and premature depolarization block may be explained largely by impaired sodium handling. However, the differences in action potential morphology observed for GABAergic neurons are more complex and suggest that depolarization block in these neurons may be due to multiple factors including impaired Na^+^ handling.

**Figure 8.**
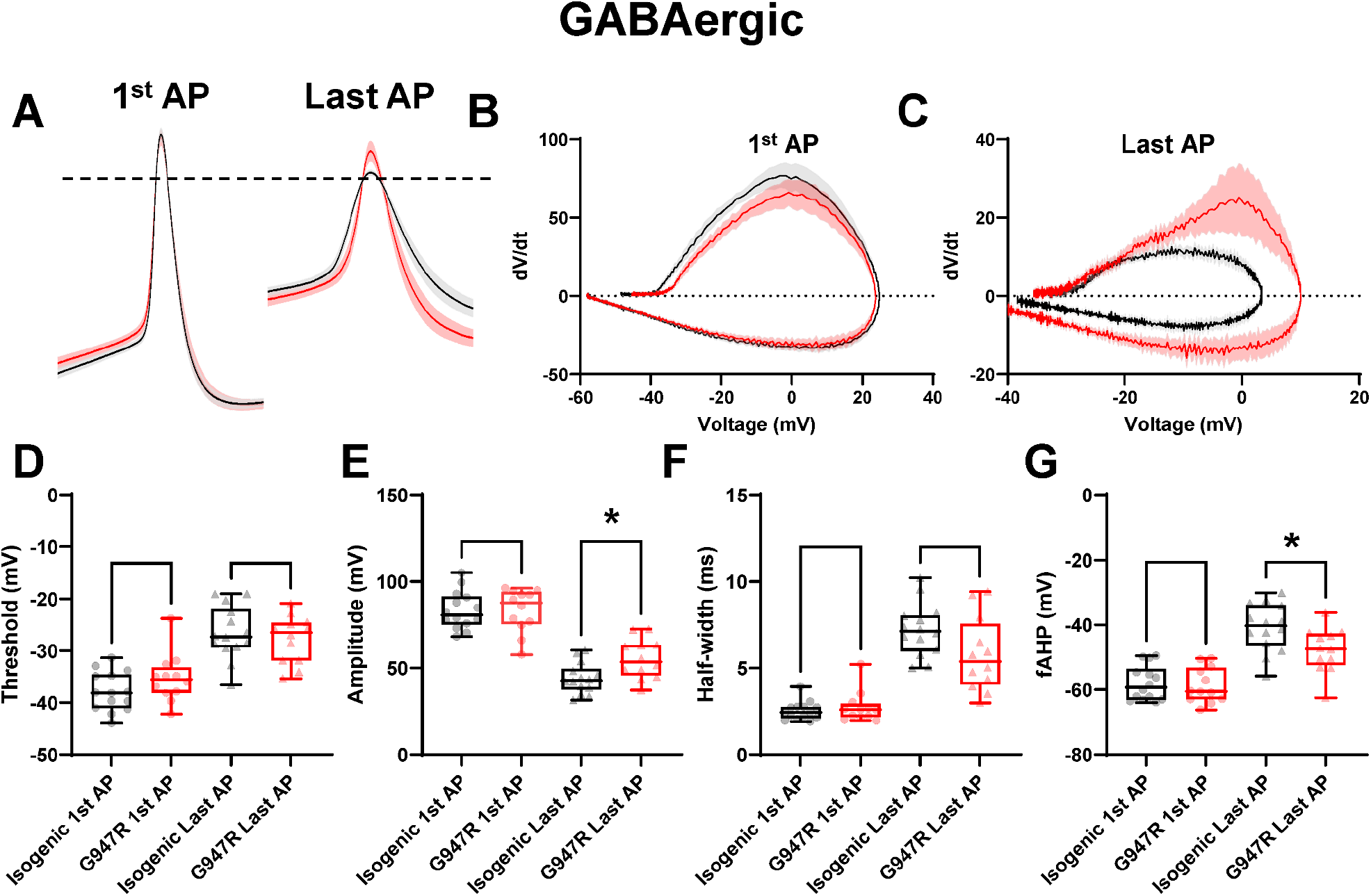
Action potential morphology of GABAergic neurons. **(A)** Averaged first and last action potentials evoked by 50 pA current inputs recorded from isogenic control (black) and G947R (red) GABAergic neurons. Averaged phase plane plots for **(B)** the first and **(C)** last action potentials evoked by 50 pA current inputs recorded from isogenic control (black) and G947R (red) GABAergic neurons. Summary data for **(D)** action potential threshold, **(E)** amplitude, **(F)** half-width, and **(G)** fAHP for the first (circles) and last (triangles) evoked by 50 pA current inputs. Average action potentials and phase plane plots are plotted as mean ± SEM and box plots represent median ± 95% CI for 19 – 23 cells. Asterisks denote p<0.05.

### AHC neurons exhibit a delayed GABA switch

In developing neurons, the neurotransmitter GABA is excitatory but becomes inhibitory later in development as expression of the sodium-potassium-chloride co-transporter NKCC1 decreases simultaneously with increased expression of the potassium chloride co-transporter KCC2, resulting in a low intracellular chloride ion concentration in mature neurons. The time at which this switch in polarity occurs is a critical developmental milestone, and some neurodevelopmental disorders exhibit a delay in this GABA switch (26–31). We measured the GABA reversal potential (*E*_GABA_) of isogenic control or G947R glutamatergic neurons over the course of 65 days to determine when the GABA polarity switch occurs. Using gramicidin perforated patch, which preserves the native chloride gradient, we sequentially applied 100 µM GABA to elicit currents at holding potentials from −80 mV to −20 mV **(Fig. 9A)** and quantified the percentage of cells for which the reversal potential was more hyperpolarized than −60 mV. Over the course of 65 days in culture, a smaller percentage of G947R neurons exhibited *E*_GABA_ values more hyperpolarized than −60mV compared with isogenic control neurons **(Fig. 9B,C)**. By 49 DIV, all isogenic control neurons had switched from excitatory to inhibitory GABA responses, while only 67% of G947R neurons exhibited the switch. Additionally, by 65 DIV, only 80% of G947R neurons showed GABA evoked inhibitory currents **(Fig. 9B,C)**, representing a substantial delay in the GABA switch for G947R neurons.

**Figure 9.**
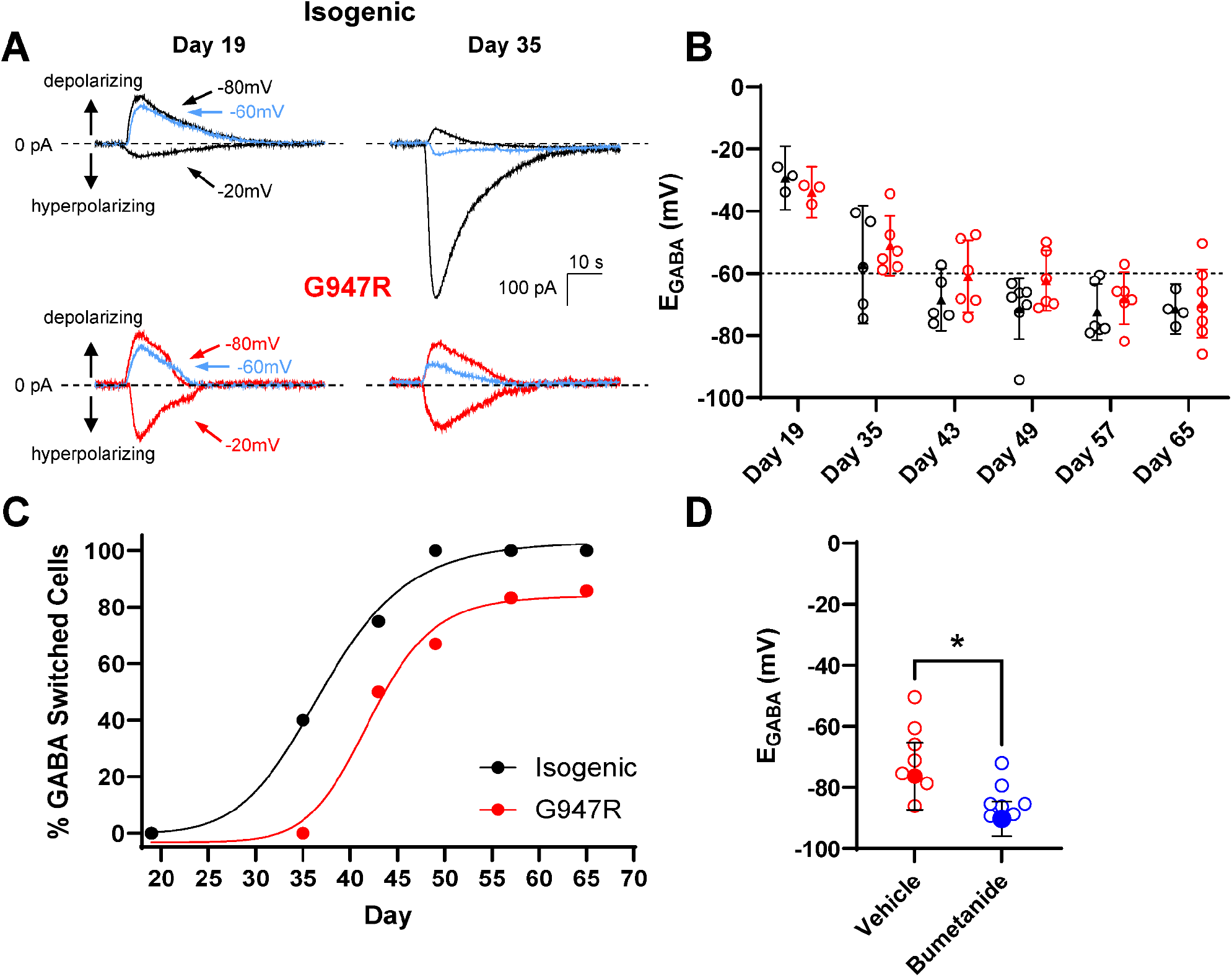
The GABA switch is delayed in ATP1A3 G947R glutamatergic neurons. **(A)** Representative GABA induced chloride currents elicited from −80 to −20 mV from isogenic control (black) and G947R glutamatergic neurons at multiple time points. The blue highlighted trace represents GABA induced chloride currents at −60mV. Current polarity is inverted for display purposes. **(B)** Summary box plots of *E*_*GABA*_ across time. Dashed line at −60 mV represents the threshold at which GABA switched from excitatory to inhibitory. **(C)** Percentage of cells in which GABA has switched from excitatory to inhibitory. **(D)** *E*_*GABA*_ of G947R neurons treated with vehicle, or 10 µM bumetanide. Box plots represent median ± 95% CI for 3 – 7 cells.

Given the importance of chloride transporters in setting *E*_GABA_, we tested whether inhibition of the Na/K/2Cl co-transporter NKCC1 would modulate the GABA response in mutant neurons. We exposed G947R neurons at 65 DIV to 10 µM bumetanide or vehicle for 2-3 hours before measuring *E*_GABA_. Bumetanide treated cells exhibited significantly more hyperpolarized *E*_GABA_ than vehicle treated cells **(Fig. 9D)**. All treated cells exhibited *E*_GABA_ more hyperpolarized than −60mV. This finding suggests that NKCC1 inhibition may be able to compensate for the delay in the GABA switch in G947R neurons.

## Discussion

Neurodevelopmental disorders (NDD) are prevalent in the general population (32) and several genetic etiologies have been identified (33–35). Investigating individual genes associated with NDD can inform about pathophysiology. In this study, we conducted an extensive investigation of the neurophysiological dysfunction of human neurons heterozygous for the recurrent *ATP1A3* variant G947R associated with AHC, a severe neurodevelopmental disorder without specific treatments. Our findings illuminate a cascade of pathophysiological events that underlie the disorder and offer new opportunities for conceiving new therapeutic strategies

The neuronal NKA encoded by *ATP1A3* is essential for establishing ionic gradients and clearance of intracellular Na^+^ following intense neuronal activity. While expression of this ion pump becomes enriched in GABAergic neurons, early in development this pump is broadly expressed across cortical regions in both glutamatergic and GABAergic neurons (20). Because AHC is a neurodevelopmental disorder, understanding the cellular mechanisms of this disease at the earliest timepoint is critical, and human iPSC-derived neurons represent an ideal model system to interrogate very early stages of neurodevelopmental diseases.

The glycine 947 residue is adjacent to the Na^+^ binding site in ATP1A3, and here we show that mutation of this residue has large effects on ion homeostasis in glutamatergic and GABAergic neurons. Our work shows that both Na^+^ and K^+^ translocation through the pump is impaired, resulting in smaller ouabain sensitive pump currents, collapsed Na^+^ and K^+^ gradients, and depolarized resting membrane potential. Other work has shown that ATP1A3-P775L variant shows an aberrant cation leak (36). However, we did not observe any difference in basal currents amplitude between ATP1A3-G947R and isogenic control neurons of either cell type that would be suggestive of aberrant ionic leak through the pump. Moreover, when using predicted intracellular K^+^ concentrations based on direct measurements of *E*_K_ as input into the Goldman-Hodgkin-Katz equation, the theoretical calculations closely align with our measured RMP values, suggesting that RMP is consistent with Nernstian predictions of ionic reversal potentials. The presence of leak Na^+^ currents would increase relative membrane Na^+^ permeability, and the effect on RMP would be greater than that predicted by the GHK equation.

Activity-dependent neuronal maturation is a complex feedforward process whereby neuronal activity leads to transcriptional changes of genes governing neuronal growth, morphology, and electrical activity in order to drive the formation of stable neuronal networks. One important aspect of this is the development of sustained neuronal firing. Here we show that both glutamatergic and GABAergic ATP1A3-G947R are more prone to depolarization block compared to isogenic control neurons. A critical component of sustained neuronal firing is clearance of Na^+^ during and immediately following intense neuronal activity, a function for which ATP1A3 is ideally suited. However, we show here that G947R glutamatergic neurons show slower recovery from high Na^+^ loads compared to isogenic control neurons. This is reflected in neuronal firing patterns as a higher propensity for depolarization block. One may speculate that inhibition of Na^+^ leak channels may be a potential therapeutic pathway.

Another major component of neuronal maturation is the timing of the GABA switch, where the activity of GABA switches from excitatory to its canonically inhibitory role in central neurons. Numerous neurodevelopmental disorders show a delay in the GABA switch (26–31), which has downstream effects on chloride homeostasis and excitation/inhibition balance. Here we show that the GABA switch is delayed in ATP1A3-G947R neurons. As a proof-of-concept, we showed that inhibition of NKCC1 by bumetanide hyperpolarized *E*_GABA_. While modulation of neuronal chloride gradients may be a potential therapeutic pathway, inhibition of NKCC1 early in development has been shown to induce deficits in excitatory neurotransmission resulting from reduced dendrite formation and dendritic spine density (37). Additionally, bumetanide is a powerful loop diuretic acting on kidney-expressed NKCC2, and systemic use may lead to severe dehydration and electrolyte imbalance. Selective targeting leading to increased KCC2 activity may be an alternative strategy (38).

In this work, we have shown that impaired pump current in both glutamatergic and GABAergic neurons lead to disrupted ionic gradients. This results in complex secondary effects on neuronal excitability, with both neuronal subtypes more prone to depolarization block, and a delayed neurodevelopmental trajectory whereby GABA does not adopt its canonical inhibitory function until later in development. Taken together, these results suggest that a profound impairment of excitatory-inhibitory balance may be a prevalent feature of AHC during very early developmental stages, where ATP1A3 plays a prominent role in neuronal physiology. How this persists throughout development as ATP1A3 becomes enriched in inhibitory neurons is an important question, and best suited to more complex model systems, including the ATP1A3-G947R mouse line (39).

## Methods

### Generation and validation of induced pluripotent stem cells

A previously reported and validated induced patient-derived pluripotent stem (iPSC) line was used for this study (25). The cell line was derived from a female diagnosed with AHC at age 11 months who experienced intermittent hemiplegic episodes and developmental delay. Genetic testing revealed the recurrent *ATP1A3* missense mutation c.2839G>A (p.G947R).

An isogenic mutation-corrected version of this iPSC line was generated using CRISPR genome editing. A DNA oligonucleotide encoding a single guide RNA was designed using CRISPOR software (http://crispor.gi.ucsc.edu/). A single stranded donor oligonucleotide was designed to enable repair of the pathogenic mutation by homology-dependent repair (HDR) (c.2839A>G) and to create synonymous single nucleotide variant (SNV) at the corrected codon (c.2841G>A) to help confirm on-target editing. The repair oligonucleotide was also designed to create a synonymous SNV at the PAM site (c.2829C>T) to limit recleavage of the target genomic DNA. The designed SNVs preserved codon optimality by having similar human codon usage frequency. Oligonucleotides were synthesized by Integrated DNA Technologies (IDT, Coralville, IA). The DNA-encoded guide RNA was inserted into plasmid pX459 encoding *S. pyogenes* Cas9 (generously provided by Feng Zhang, the Broad Institute, Cambridge, MA; Addgene Catalog #62988) at the BbsI site and verified by Sanger sequencing. The previously reported AHC patient-derived iPSC line was electroporated with the plasmid and HDR oligonucleotide using a MaxCyte STX instrument. After puromycin selection for 48 h and continued culture for 2 weeks in a hypoxic atmosphere (5% CO_2_, 5% O_2_, 37°C), single colonies were manually selected, re-plated, expanded, then genotyped by PCR and Sanger sequencing. One identified iPSC clone had the corrected mutant base pair, no on-target editing errors, and was heterozygous for synonymous SNVs at both the corrected codon-947 and PAM site (**Fig. 1**). The detection of heterozygous SNVs excludes monoallelic amplification of the WT allele (40). The mutation-corrected isogenic iPSC line was expanded and evaluated for off-target edits using PCR-sequencing. There were no unintended edits for predicted genomic regions with 3 or fewer mismatches to the guide RNA. Chromosomal integrity was assessed using G-band karyotyping (Cell Line Genetics, Madison, WI). Cytogenetic analysis demonstrated a normal female karyotype (**Supplemental Figure S1**).

### Differentiation of human iPSC-neurons

Human iPSC-derived cortical excitatory glutamatergic neurons were generated by neurogenin-2 (NGN2) induction with modifications.(41, 42) Briefly, human iPSCs in suspension were transduced by separate lentiviral vectors encoding NGN2 driven by a tetracycline-inducible promoter (TetO-NGN2), reverse tetracycline-controlled transactivator (rtTA) and red fluorescent protein (RFP) then plated on matrigel-coated plates in E8 medium (43) containing 10 µM Y27632 (StemCell technologies). Cells were maintained in a transitional medium from knockout serum replacement to neural induction medium with the supplements LDN-193189 (100 nM, Stemgent, Lexington, MA), SB431542 (10 µM, StemGent), and XAV939 (2 µM, Sigma-Aldrich, St. Louis, MO). NGN2 and RFP expression were induced by 2 µg/ml doxycycline one day prior to a 48-hour selection in puromycin (2 µg/ml). After day 5, induced neuronal cells were plated on primary mouse glial cells cultured on poly-D-lysine/laminin coated coverslips, and continually maintained in Neurobasal medium supplemented with N2, B27, BDNF (10 ng/ml, Pepro-Tech, Rocky Hill, NJ) and doxycycline (2 µg/ml) for up to 35 days. AraC (4 µM) was added on day 7 through 12. Differentiated neurons were identified by morphology and red fluorescence then used for electrophysiology studies.

Human iPSC-derived cortical GABAergic (iGABA) neurons were generated by induction with ASCL1 and DLX2 with modifications to the reported protocol (44). iGABA neurons were generated by transduction with three separate doxycycline-inducible lentiviral vectors encoding tetO-ASCL1-PuroR, tetO-DLX2-HygroR, and green fluorescent protein (GFP) (44). Briefly, iPSCs were harvested in Accutase, dissociated into a single-cell solution, quenched in DMEM, pelleted by centrifugation for 5 min at 800 × g, and resuspended in mTeSR with 10 μM ROCK Inhibitor. Volumetric equivalents of tetO-ASCL1-PuroR, tetO-DLX2-HygroR, and pUBIQ-rtTA were added to the suspension, mixed gently by inversion, dispensed onto Matrigel-coated plates, and incubated overnight at 37°C. The next day, media was changed to Induction Media with 3 μg/mL doxycycline (DIV1). 2 μg/mL puromycin and 25 μg/mL hygromycin (Thermo, #10687010) were added the next day (DIV2) and continued for four days. On day 5, induced neuronal cells were plated on primary mouse glial cells cultured on poly-D-lysine/laminin coated coverslips, and continually maintained in Neurobasal medium supplemented with N2, B27, BDNF (10 ng/ml, PeproTech, Rocky Hill, NJ) and doxycycline (2 µg/ml) for up to 65 days. AraC (4.0 µM) was added on day 7 through 12. Differentiated neurons were identified initially by morphology and then secondarily by red fluorescence for glutamatergic neurons and green fluorescence for GABAergic neurons.

Co-cultures of cortical iGABA and excitatory neurons were done by plating cells in a ratio of 30:70 on primary mouse glial cells and cultured for up to 65 days.

### Measurement of pump current

Pump current was recorded by whole-cell voltage-clamp at room temperature using an Axopatch 200B amplifier (Molecular Devices, San Jose, CA USA). Patch pipettes were pulled from borosilicate glass capillaries (Warner Instruments, Hamden, CT, USA) with a multistage P-1000 Flaming-Brown micropipette puller (Sutter Instruments Co., San Rafael, CA, USA) and fire-polished using a microforge (Narashige MF-830; Tokyo, Japan) to a resistance of 1.5–2.5 MΩ. For measurement of pump current, the pipette solution contained (in mM) 95 NaMeSO_3_, 20 TEA-Cl, 2 BaCl_2_, 5 EGTA, 4 MgATP, 5 Tris-phosphocreatine, 10 HEPES. 5 glucose at pH 7.35 (adjusted with NaOH) and osmolality 280 mOsmol/kg (adjusted with sucrose). Initially, cells were continuously superfused with a K^+^-free recording solution containing (in mM) 140 NaCl, 1 MgCl_2_, 2 BaCl_2_, 2 CsOH, 2 NiCl_2_, 0.2 CdCl_2_, 0.33 NaH_2_PO_4_, 5 HEPES. 5 glucose, 20 NMDG-Cl at pH 7.35 (adjusted with HCl). Pump currents were activated by superfusing cells with an extracellular solution in which 20 mM NMDG-Cl was replaced with 20 mM KCl. Cells were held at +40 mV for the duration of recordings. Pump current was measured sequentially in the absence and presence of 10 µM ouabain. Ouabain was prepared as 10 mM stock in bath solution and diluted to 10 μM in extracellular solution on the day of experiments. Ouabain-sensitive current was calculated by subtracting ouabain-resistant current from total current measured initially in the absence of ouabain. Pump current measurements were performed at 34-36 days in vitro (DIV) for induced glutamatergic neurons and GABAergic neurons co-cultured with glutamatergic neurons were assayed at 60-63 DIV.

### Measurement of potassium equilibrium potential

For measurement of potassium equilibrium potential (*E*_K_), voltage-clamp recordings were made in cell-attached mode at room temperature using pipettes with resistances of 1.0-1.2 MΩ. The pipette solution consisted of (in mM): 155 NaCl, 3.5 KCl, 1 MgCl_2_, 1.5 CaCl_2_, 2 NiCl_2_, 0.2 CdCl_2_, 10 HEPES, 10 glucose with pH 7.35. Bath solution consisted of (in mM) 155 NaCl, 3.5 KCl, 1 MgCl_2_, 1.5 CaCl_2_, 10 HEPES, 10 glucose with pH 7.35. Equilibrium potential was measured during voltage ramps from ™120 to +80 mV over 500 msec, and *E*_K_ was calculated as the x-intercept for the evoked current. Intracellular K^+^ concentration, and theoretical membrane potential were calculated using the Nernst equation and Goldman-Hodgkin-Katz equation, respectively (www.physiologyweb.com/calculators). Measurements were performed at 34-36 DIV for glutamatergic neurons and GABAergic neurons co-cultured with glutamatergic neurons were assayed at 60-63 DIV.

### Perforated patch current clamp recordings

Action potentials were recorded from neuronal cell bodies by whole-cell current clamp using a MultiClamp 700B amplifier (Molecular Devices, San Jose, CA, USA). Patch pipettes were fabricated from borosilicate glass (World Precision Instruments, Sarasota, FL, USA) to have resistance 3.0-4.0 MΩ when filled with a solution containing (in mM) 110 K-gluconate, 10 NaCl, 10 KCl, 0.1 CaCl_2,_ 10 HEPES, 5 EGTA, 10 trisphosphocreatine, 4 Mg-ATP, 0.3 Na-GTP with pH 7.35 and osmolality 280 mOsmol/kg. Bath solution consisted of (in mM) 155 NaCl, 3.5 KCl, 1 MgCl_2_, 1.5 CaCl_2_, 10 HEPES, 10 glucose with pH 7.35.

We used amphotericin B perforated patch to assess susceptibility of neurons to depolarization block. Amphotericin B was resuspended in DMSO at a stock concentration of 60 mg/mL. On the day of recording experiments, amphotericin B was diluted to a working concentration of 200 µg/mL in the same pipette solution used for current clamp recording. Solutions were sonicated and vortexed vigorous to promote dissolution of the ionophore. Amphotericin B containing internal solution was protected from light and replaced every 3 hours. After gigaohm seal formation, cells were clamped at −80 mV and access resistance (Ra) was continuously monitored. Current clamp experiments commenced after Ra fell to 20 MΩ or less (indicating formation of the perforated patch) and resting membrane potential (RMP) was measured immediately. To assess evoked action potential activity and depolarization block, we used both step and ramp protocols. Action potentials were evoked by 2-s steps from −30 to 210 pA from a holding potential of −65 mV. Action potentials were also evoked by a 3-s current injection ramp from −30 pA to 500 pA. The ramp protocol was run in triplicate and the average of the three trials was used for analysis. Cells for which the perforated patch ruptured into whole-cell configuration were excluded from analysis. Measurements for glutamatergic neurons were performed at 34-36 DIV and GABAergic neurons co-cultured with glutamatergic neurons were assayed at 60-63 DIV.

Evoked action potential firing and morphology were assessed using a custom MatLab script (available at https://prism.northwestern.edu/records/zpfr2-ysm22).Only action potentials more depolarized than 0 mV with an upstroke velocity of 5 mV/ms were counted. Action potential threshold was calculated as the maximum value of the second derivative of the action potential waveform, action potential amplitude was measured as the maximal voltage during the action potential waveform, and action potential width was measured at half-maximal spike amplitude. Fast afterhyperpolarization (fAHP) was measured as the most negative absolute voltage following an action potential. Time to depolarization block was defined as the time from the beginning of the stimulus to the cessation of repetitive firing and stabilization of a depolarized membrane potential. The number of action potentials fired and current injection required to induce depolarization block were assessed using a ramp protocol. The number of action potentials fired was counted from the first to the last action potentials with peak depolarization more positive than 0 mV. Current magnitude required to induce depolarization block was assessed as the current required to induce a stable depolarized membrane potential.

### Determination of GABA equilibrium potential (*E*_GABA_)

*E*_GABA_ was measured in glutamatergic neurons between 19-65 DIV using gramicidin perforated patch clamp recording. Gramicidin was resuspended in DMSO at a concentration of 10 mg/mL, and stock solutions were stored at −20°C. On the day of recording, gramicidin was diluted to a working concentration of 25 µg/mL in pipette solution containing (in mM): 145 KCl, 5 EGTA, 10 HEPES (pH 7.2 adjusted with KOH) by sonication and vigorous vortexing. Gramicidin containing internal solution was kept protected from light, and replaced every 3 hours. Extracellular solution was the same as described for current clamp recording. Following gigaohm seal formation, cells were clamped at −80 mV and Ra was continuously monitored. *E*_GABA_ was measured by voltage clamp after Ra fell to ≤ 20 MΩ. GABA receptor mediated chloride currents were elicited by a 10 s application of 100 µM GABA, with 3 minutes between each application. Cells were clamped from −90 to −20 in 10 mV increments. To test whether bumetanide was able to induce hyperpolarization of *E*_*GABA*_, G947R cells were treated with 10 µM bumetanide for 2-3 hours prior to recording. *E*_GABA_ was determined as the x-intercept from a linear regression of the current-voltage relationship of GABA-evoked currents. Cells for which the perforated patch ruptured into whole-cell configuration were excluded from analysis.

### Sodium ion imaging

We used the Na^+^ selective dye ION-Natrium Green 2-AM (ING-2 AM, ION Bioscience, San Marcos, TX) to measure basal Na^+^ concentration and the kinetics of intracellular Na^+^ change after Na^+^ loading. ING-2 AM was resuspended in DMSO at a stock concentration of 46 mM, and diluted to 4.6 µM in dye loading solution containing (in mM): 10 NaCl, 100 Tris-HCl, 5.4 KCl, 1.3 CaCl_2_, 0.8 MgSO_4_, 0.78 NaH_2_PO_4_, 10 HEPES, 20 glucose, 1% bovine serum albumin (BSA) with pH 7.3, and osmolality 320 mOsm/kg. Glutamatergic neurons were loaded with ING-2 AM for 30 minutes at 37°C, then washed 3 times with a low Na^+^ solution having the same composition as the dye loading solution but without BSA. Cells were then loaded into an imaging chamber and continuously perfused with low Na^+^ solution bubbled with 95% O_2_ / 5% CO_2_ at 35 ± 1.0°C. Basal conditions were recorded for 5 minutes then cells were Na^+^ loaded by superfusing 5 minutes with a high Na^+^ solution containing solution (in mM) 155 NaCl, 3.5 KCl, 1 MgCl_2_, 1.5 CaCl_2_, 10 HEPES, 10 glucose with pH 8.2 to achieve a final pH of 7.3 resulting from acidification by bubbling with 95% O_2_ / 5% CO_2_. Cells were then switched back to low Na^+^ solution and monitored over time. Images were captured using a Hamamatsu OrcaFlash 4.0LT camera every 5 seconds at 20X magnification. ING-2 fluorescence was excited using a standard EGFP filter set (Chroma 49002, Bellows Falls, VT) using an XCite 110LED illumination system (Excelitas, Pittsburgh, PA).

Automated image segmentation was performed using Cellpose-SAM (45), and regions of interest (ROI) were exported for fluorescence intensity analysis using FIJI (46). All images were background-subtracted and corrected for photobleaching. Basal ING-2 fluorescence was determined as the average background subtracted, photobleaching-corrected fluorescence intensity during the first minute of recording under low Na^+^ conditions and expressed as arbitrary units (A.U.). Sodium recovery was measured as the time constant of the recovery phase following Na^+^ loading. All time constants were fit using an exponential function in Clampfit. Cells for which an exponential function could not be fit were excluded from analysis. Sodium imaging experiments were performed on glutamatergic neurons at 35-40 DIV.

### Data analysis

Electrophysiological data were analyzed using a combination of Clampfit

10.4 (Molecular Devices), Excel 2019 (Microsoft, Redmond, WA, USA), GraphPad Prism (GraphPad Software, San Deigo, CA, USA), and custom MatLab scripts. Results are presented as mean ± 95% confidence interval (CI) unless otherwise noted. For electrophysiology data, values more than two standard deviations from the mean were considered outliers and excluded from analyses, which resulted in the exclusion of less than 2.5% of individual cell recordings. Statistical comparisons were performed using two-way ANOVA or unpaired two-tailed Student’s *t*-test when appropriate, and p-values < 0.05 were considered statistically significant.

## Acknowledgements

We thank Jean-Marc DeKeyser for technical assistance.

## Funding

This work was supported by National Institutes of Health grant NS125785.

## Notes

### Competing Interest Statement

The authors have declared no competing interest.

