## Supplementary Information for "ATP1A3 dysfunction alters excitability and developmental trajectory of human neurons in alternating hemiplegia of childhood"

### Supplementary Figures

**Figure S1.** Cytogenetic analysis of isogenic control iPSC line

**Figure S2**. Current required to induce depolarization block in ATP1A3-G947R neurons

**Figure S3.** Action potential upstroke and downstroke velocity in ATP1A3-G947R neurons

**Figure S4.** Input resistance of ATP1A3-G947R neurons

**Figure S5.** Baseline currents at +40 mV of ATP1A3-G947R neurons

### Supplementary Tables

**Table S1.** Pump current measurements in iPSC-neurons

**Table S2.** Properties of ionic gradients in iPSC-neurons

**Table S3.** Depolarization block in iPSC-neurons

**Table S4.** Action potential morphology of glutamatergic neurons

**Table S5.** Action potential morphology of GABAergic neurons

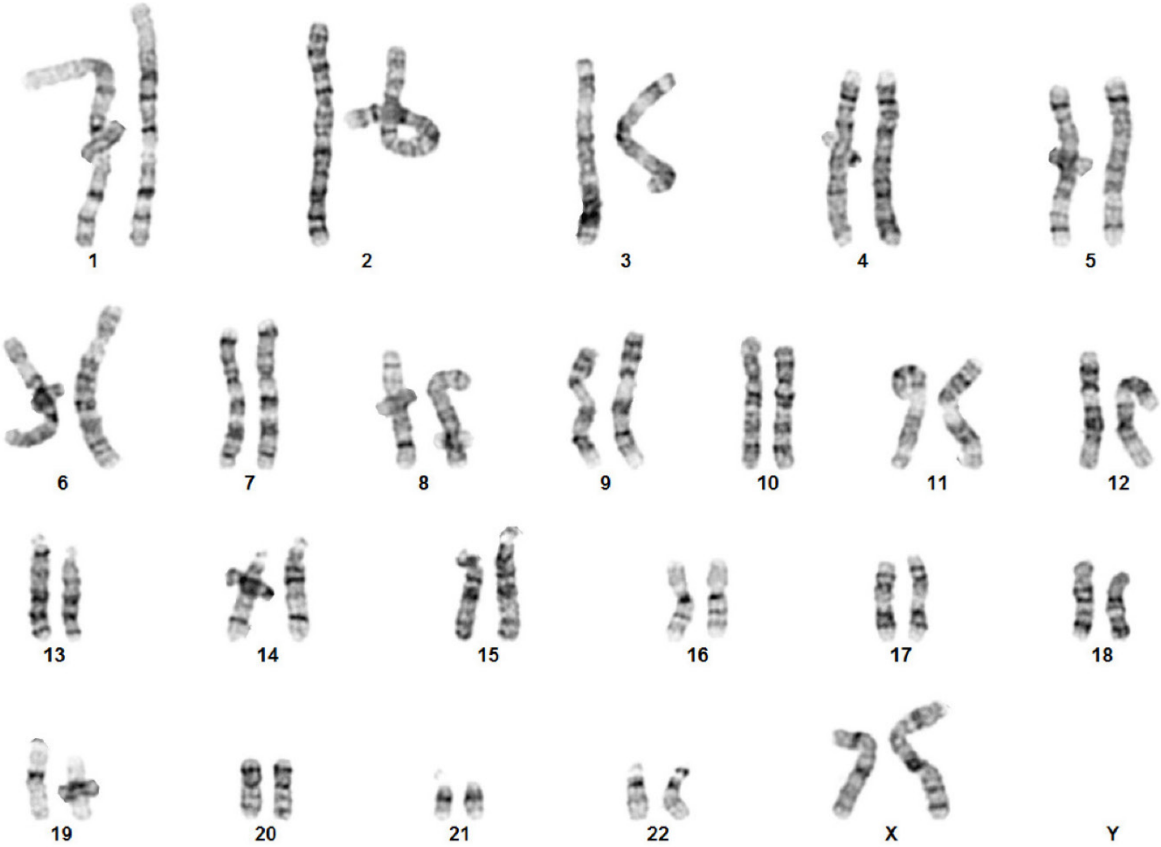


**Figure S1. Cytogenetic analysis of isogenic control iPSC line**

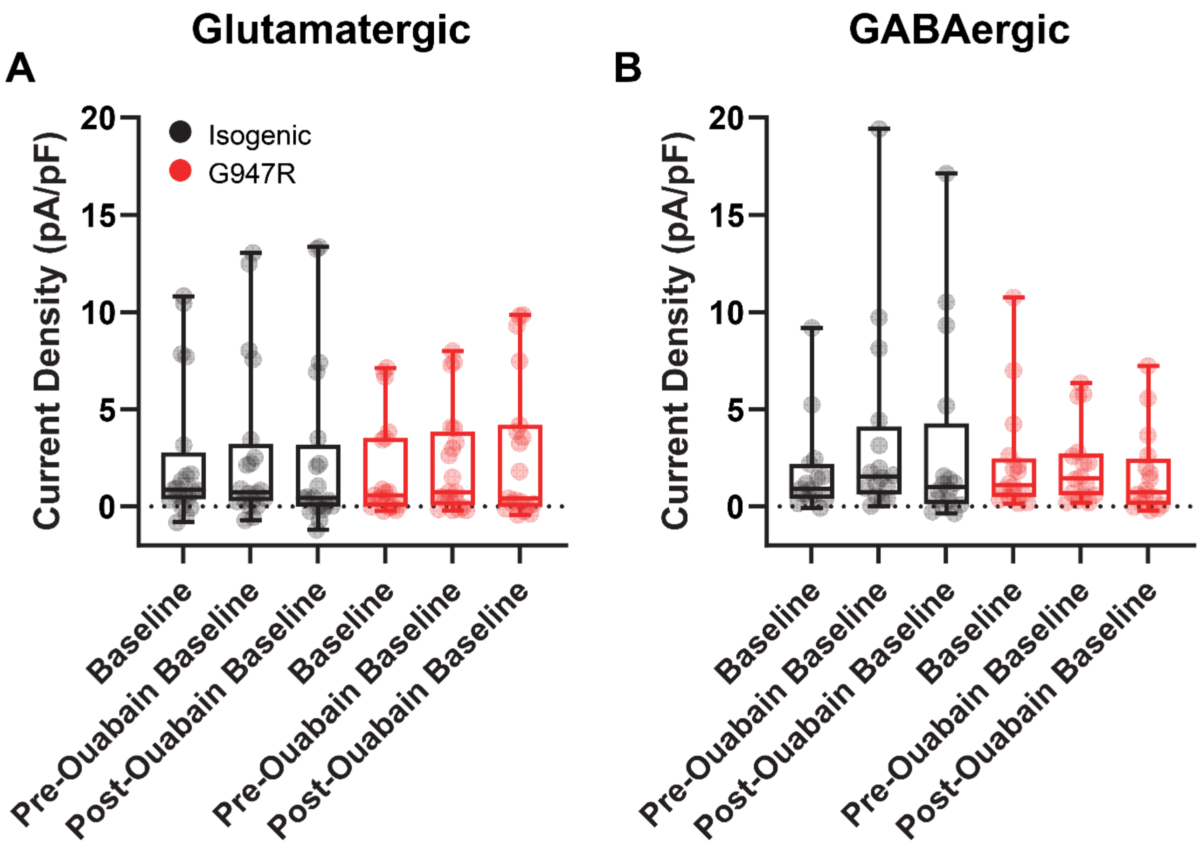


**Figure S2. Baseline currents at +40 mV of ATP1A3-G947R neurons.** Summary data showing baseline currents at +40 mV prior to the first pump activation, prior to ouabain application, and prior to the second pump activation for (A) glutamatergic and (B) GABAergic neurons. Box plots represent median ± 95% CI for 15 – 20 cells. Asterisks denote $p<0.05$.

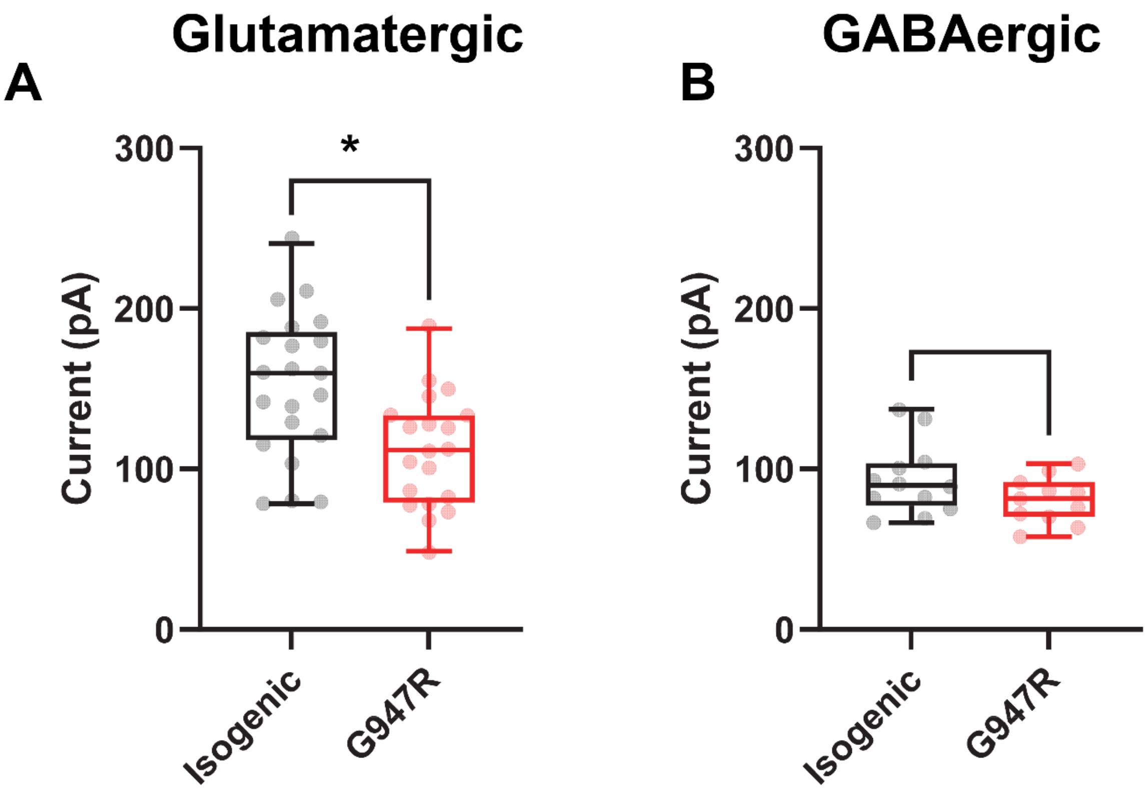


**Figure S3. Current required to induce depolarization block in ATP1A3-G947R neurons.** Summary data showing current required to induce depolarization block elicited by a ramp current injection for (A) glutamatergic and (B) GABAergic neurons. Box plots represent median ± 95% CI for 11 – 21 cells. Asterisks denote $p<0.05$.

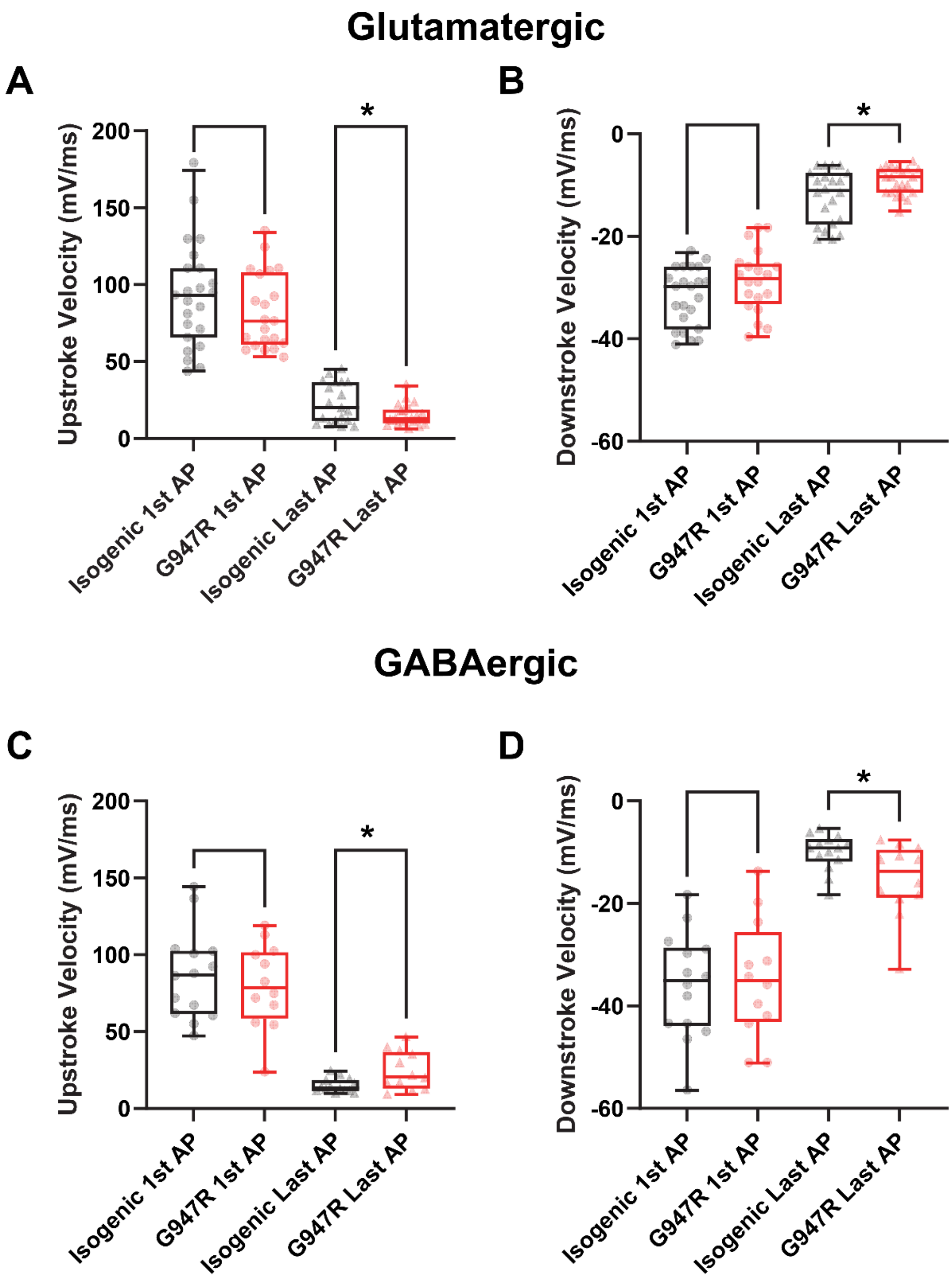


**Figure S4. Action potential upstroke and downstroke velocity in ATP1A3-G947R neurons.** Summary data showing **(A)** upstroke and **(B)** downstroke velocity of the first and last action potentials in a train elicited by a 100 pA current injection in glutamatergic neurons. Summary data showing **(C)** upstroke and **(D)** downstroke velocity of the first and last action potentials in a train elicited by a 50 pA current injection in glutamatergic neurons. Box plots represent median ± 95% CI for 12 – 23 cells. Asterisks denote $p<0.05$.

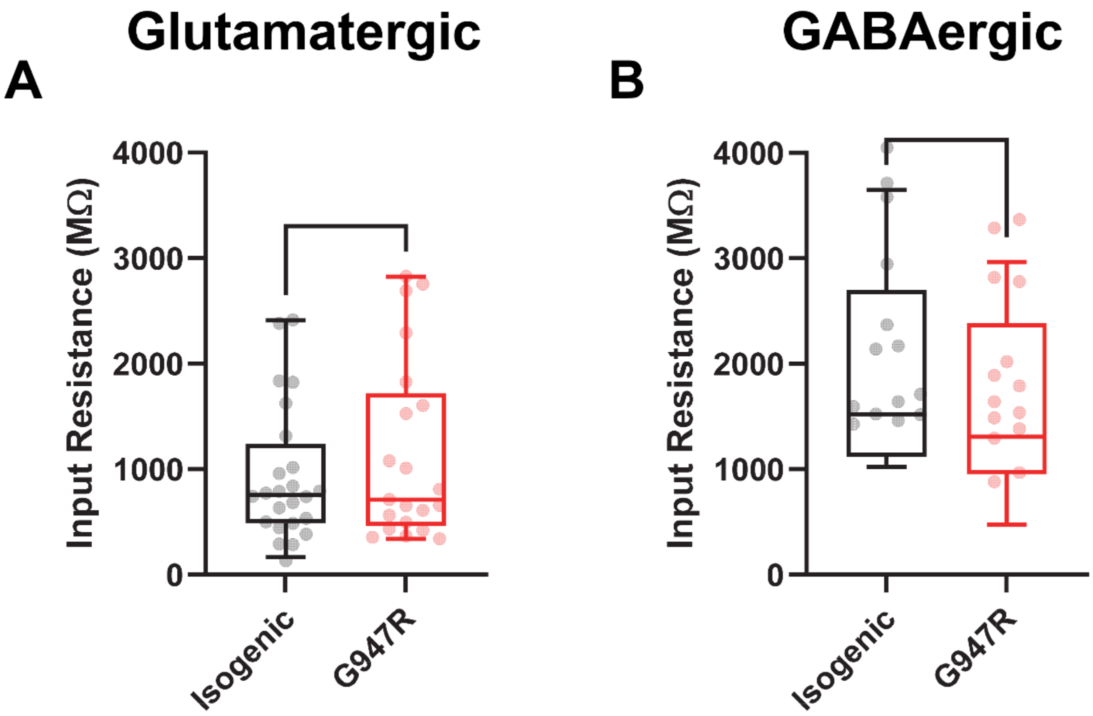


**Figure S5. Input resistance of ATP1A3-G947R neurons.** Summary showing input resistance of (A) glutamatergic and (B) GABAergic neurons. Box plots represent median ± 95% CI for 14 – 24 cells. Asterisks denote $p<0.05$.

| Supplemental Table 1: Pump current measurements in iPSC-neurons | | | |
|---|---|---|---|
| *Glutamatergic neurons* | **Isogenic** | **G947R** | **p-value** |
| **Total pump current (pA/pF)** Mean ± 95% CI | 1.2 ± 0.2 (22)* | 0.8 ± 0.1 (21) | 0.000114 |
| **Ouabain-sensitive pump current (pA/pF)** Mean ± 95% CI | 0.8 ± 0.1 (22) | 0.3 ± 0.1 (21) | 0.000001 |
| **Average baseline current density (pA/pF)** Mean ± 95% CI | 2.4 ± 1.6 (20) | 1.9 ± 1.2 (19) | 0.969116‡ |
| **Average baseline + Pre-Ouabain current density (pA/pF)** Mean ± 95% CI | 2.7 ± 1.8 (20) | 2.2 ± 1.2 (20) | 0.969116‡ |
| **Average baseline + Post-Ouabain current density (pA/pF)** Mean ± 95% CI | 2.5 ± 1.9 (20) | 2.8 ± 1.7 (19) | 0.969116‡ |
| *GABAergic neurons* | | | |
| **Total pump current (pA/pF)** Mean ± 95% CI | 1.2 ± 0.2 (17) | 0.8 ± 0.2 (17) | 0.002882 |
| **Ouabain-sensitive pump current (pA/pF)** Mean ± 95% CI | 0.7 ± 0.2 (17) | 0.4 ± 0.1 (17) | 0.010748 |
| **Average baseline current density (pA/pF)** Mean ± 95% CI | 1.8 ± 1.2 (15) | 2.3 ± 1.4 (16) | 0.642251‡ |
| **Average baseline + Pre-Ouabain current density (pA/pF)** Mean ± 95% CI | 3.5 ± 2.5 (16) | 2.1 ± 1 (16) | 0.642251‡ |
| **Average baseline + Post-Ouabain current density (pA/pF)** Mean ± 95% CI | 3.1 ± 2.5 (16) | 1.7 ± 1.1 (16) | 0.642251‡ |

`

*Values in parenthesis represent the number of cells assayed

‡ p-value from One-way ANOVA comparing Average baseline current density and Average baseline + Ouabain current density between genotypes

| **Supplemental Table 2: Properties of ionic gradients in iPSC-neurons** | | | |
|---|---|---|---|
| ***Glutamatergic neurons*** | **Isogenic** | **G947R** | **p-value** |
| **$E_K$ (mV)**<br>Mean ± 95% CI | -93.6 ± 9.1 (7)* | -68.5 ± 12.0 (9) | 0.008027 |
| **Intracellular $K^+$ concentration (mM)**<br>Mean ± 95% CI | 116.7 ± 28.0 (6) | 53.1 ± 24.5 (8) | 0.005842 |
| **RMP (mV)**<br>Mean ± 95% CI | -55.6 ± 2.3 (17) | -47.9 ± 3.1 (21) | 0.000713 |
| **Basal ING-2 fluorescence (AU)**<br>Mean ± 95% CI | 181.6 ± 10 (253) | 222.5 ± 9.3 (321) | <0.000001 |
| **$Na^+$ recovery time constant (ms)**<br>Mean ± 95% CI | 0.65 ± 0.03 (250) | 0.96 ± 0.03 (304) | <0.000001 |
| ***GABAergic neurons*** | | | |
| **$E_K$ (mV)**<br>Mean ± 95% CI | -92.3 ± 4.7 (14) | -80.9 ± 6.4 (11) | 0.008484 |
| **Intracellular $K^+$ concentration (mM)**<br>Mean ± 95% CI | 131.9 ± 26.7 (14) | 86.0 ± 19.3 (11) | 0.016187 |
| **RMP (mV)**<br>Mean ± 95% CI | -61.1 ± 1.6 (9) | -55.5 ± 2.1 (12) | 0.000879 |

*Values in parenthesis represent the number of cells assayed

**Supplemental Table 3: Depolarization block in iPSC-neurons**

| *Glutamatergic neurons* | **Isogenic** | **G947R** | **p-value** |
|---|---|---|---|
| **Time to depolarization block (s)** Mean ± 95% CI | 1.2 ± 0.2 (21)* | 0.9 ± 0.1 (21) | 0.000905 |
| **Action potentials elicited from ramp (#)** Mean ± 95% CI | 9.6 ± 2.6 (21) | 5.0 ± 1.0 (19) | 0.003616 |
| **Maximum depolarization (mV)** Mean ± 95% CI | -19.0 ± 2.7 (22) | -24.5 ± 2.3 (20) | 0.004004 |
| **Current required to induce block (pA)** Mean ± 95% CI | 152.2 ± 19.5 (21) | 111.4 ± 15.4 (20) | 0.002757 |
| *GABAergic neurons* | | | |
| **Time to depolarization block (s)** Mean ± 95% CI | 1.0 ± 0.3 (12) | 0.6 ± 0.2 (13) | 0.024119 |
| **Action potentials elicited from ramp (#)** Mean ± 95% CI | 4.7 ± 0.8 (12) | 3.0 ± 1 (11) | 0.017907 |
| **Maximum depolarization (mV)** Mean ± 95% CI | -21.7 ± 2.7 (12) | -30.4 ± 2.9 (12) | 0.000273 |
| **Current required to induce block (pA)** Mean ± 95% CI | 93.6 ± 12.6 (12) | 80.8 ± 8.5 (11) | 0.118317 |

*Values in parenthesis represent the number of cells assayed

| Supplemental Table 4: Action potential morphology of glutamatergic neurons | | | |
|---|---|---|---|
| | **Isogenic** | **G947R** | **p-value** |
| **AP threshold (mV)** | | | |
| **1st AP**<br>Mean ± 95% CI | -38.3 ± 2.6 (23)* | -39.7 ± 2 (20) | 0.410268 |
| **Last AP**<br>Mean ± 95% CI | -27.4 ± 2.5 (20) | -22.6 ± 2.7 (20) | 0.016091 |
| **AP upstroke velocity (dV/dt)** | | | |
| **1st AP**<br>Mean ± 95% CI | 92.9 ± 14.1 (23) | 82.4 ± 10.6 (21) | 0.256192 |
| **Last AP**<br>Mean ± 95% CI | 23.1 ± 5.9 (19) | 15.1 ± 3 (21) | 0.01937 |
| **AP downstroke velocity (dV/dt)** | | | |
| **1st AP**<br>Mean ± 95% CI | -31.9 ± 2.3 (23) | -28.7 ± 2.6 (22) | 0.086943 |
| **Last AP**<br>Mean ± 95% CI | -12.2 ± 2.3 (20) | -9.3 ± 1.2 (21) | 0.025013 |
| **AP amplitude (mV)** | | | |
| **1st AP**<br>Mean ± 95% CI | 85.9 ± 4 (23) | 79.3 ± 3.9 (22) | 0.027111 |
| **Last AP**<br>Mean ± 95% CI | 52.1 ± 7.5 (21) | 42.0 ± 3.8 (21) | 0.022485 |
| **AP half-width (ms)** | | | |
| **1st AP**<br>Mean ± 95% CI | 2.7 ± 0.1 (23) | 2.7 ± 0.2 (20) | 0.956049 |
| **Last AP**<br>Mean ± 95% CI | 6.3 ± 0.8 (19) | 7.6 ± 0.8 (21) | 0.037169 |
| **fAHP (mV)** | | | |
| **1st AP**<br>Mean ± 95% CI | -53.7 ± 2.1 (23) | -49.9 ± 2.1 (20) | 0.018798 |
| **Last AP**<br>Mean ± 95% CI | -40.3 ± 3.4 (20) | -35.3 ± 2.6 (21) | 0.026479 |
| **Input resistance (MΩ)**<br>Mean ± 95% CI | 933.2 ± 256.4 (24) | 1144.0 ± 367.9 (21) | 0.353082 |

*Values in parenthesis represent the number of cells assayed

**Supplemental Table 5: Action potential morphology of GABAergic neurons**

| | Isogenic | G947R | p-value |
|---|---|---|---|
| **AP threshold (mV)** | | | |
| **1st AP**<br>Mean ± 95% CI | -37.9 ± 2 (14)* | -35.3 ± 2.4 (14) | 0.138375 |
| **Last AP**<br>Mean ± 95% CI | -26.6 ± 2.9 (12) | -27.8 ± 2.6 (12) | 0.521477 |
| **AP upstroke velocity (dV/dt)** | | | |
| **1st AP**<br>Mean ± 95% CI | 86.9 ± 15.3 (14) | 79.8 ± 14.5 (14) | 0.530171 |
| **Last AP**<br>Mean ± 95% CI | 14.8 ± 2.6 (12) | 24.6 ± 7.1 (12) | 0.011681 |
| **AP downstroke velocity (dV/dt)** | | | |
| **1st AP**<br>Mean ± 95% CI | -36 ± 5.4 (14) | -34.8 ± 6.1 (14) | 0.78638 |
| **Last AP**<br>Mean ± 95% CI | -10.0 ± 2 (12) | -15.4 ± 4.1 (12) | 0.020961 |
| **AP amplitude (mV)** | | | |
| **1st AP**<br>Mean ± 95% CI | 83.3 ± 5.7 (14) | 83.3 ± 6.6 (14) | 0.992201 |
| **Last AP**<br>Mean ± 95% CI | 44.0 ± 5 (12) | 55.3 ± 6.5 (12) | 0.009115 |
| **AP half-width (ms)** | | | |
| **1st AP**<br>Mean ± 95% CI | 2.5 ± 0.3 (14) | 2.7 ± 0.5 (14) | 0.461511 |
| **Last AP**<br>Mean ± 95% CI | 7.2 ± 0.9 (12) | 5.8 ± 1.2 (12) | 0.059719 |
| **fAHP (mV)** | | | |
| **1st AP**<br>Mean ± 95% CI | -58.1 ± 2.8 (14) | -58.7 ± 2.8 (14) | 0.783667 |
| **Last AP**<br>Mean ± 95% CI | -40.9 ± 4.2 (12) | -47.8 ± 3.9 (12) | 0.021521 |
| **Input resistance (MΩ)**<br>Mean ± 95% CI | 1870.0 ± 485.3 (14) | 1534.0 ± 426.8 (14) | 0.317424 |

*Values in parenthesis represent the number of cells assayed
